# Collective dynamics of formin and microtubule and its crosstalk mediated by FHDC1

**DOI:** 10.1101/2023.07.25.550329

**Authors:** Chee San Tong, Maohan Su, He Sun, Xiang Le Chua, Su Guo, Ravinraj S/O Ramaraj, Ong Nicole Wen Pei, Ann Gie Lee, Yansong Miao, Min Wu

## Abstract

The coordination between actin and microtubule network is crucial, yet our understanding of the underlying mechanisms remains limited. In this study, we used travelling waves in the cell cortex to characterize the collective dynamics of cytoskeletal networks. Our findings show that Cdc42 and F-BAR-dependent actin waves in mast cells are mainly driven by formin-mediated actin polymerization, with the microtubule-binding formin FH2 domain-containing protein 1 (FHDC1) identified as an early regulator. The depolymerization of microtubules coincides with the nucleation of actin waves, and the concurrent release of FHDC1 from microtubule is required for actin waves. Lastly, we show the importance of the actin-microtubule linkage mediated by FHDC1 in crucial cellular processes such as cell division and migration. Our data provided molecular insights into the nucleation mechanisms of actin waves and uncover an antagonistic interplay between microtubule and actin polymerization in their collective dynamics.

## Introduction

Actin filaments and microtubules play integral roles in virtually all cellular processes, including cell motility, division, and trafficking. Both actin filaments and microtubules are highly dynamic and undergo continuous growth and shrinkage to facilitate rapid cellular responses to environment cues. In addition, it has become interestingly clear that the interaction and coordination between actin and microtubule cytoskeletal networks are crucial for a wide array of cellular activities (Rodriguez et al., 2003; Akhshi et al., 2014).

Significant advancements have been made in understanding the mechanistic interactions between actin and microtubules through *in vitro* (Griffith and Pollard, 1978) and cell-free (Sider et al., 1999; Waterman-storer et al., 2000; Colin et al., 2018) biochemical systems over the past four decades. These reconstitution approaches have been proven valuable in dissecting the effects of proteins operating at the interface between actin and microtubule on nucleation (Okada et al., 2010; Henty-Ridilla et al., 2016), stability (Bartolini et al., 2008) and co-organization of these filaments (Sattilaro, 1986; Roger et al., 2004; Preciado López et al., 2014; Elie et al., 2015). Yet, elucidating the cooperative behavior and mutual influence of cytoskeletal components in living cells still presents considerable challenges. The complexity arises from factors such as the crowdedness of cytoskeletal network, the complex trajectories and dynamics of the cytoskeleton, and higher-order feedback loops that exist between them in a living system. Some of the useful systems used in the literature to elucidate the crosstalk between actin and microtubules in cells are the focal adhesion (Palazzo and Gundersen, 2002; Stehbens and Wittmann, 2012) or the adherent junction (Higashi et al., 2018), both of which could provide localized signals that facilitate the analysis. Recently, actin waves have emerged as a common theme of cortical organization in many single cell systems (Roy, 2016; Beta and Kruse, 2017; Devreotes et al., 2017; Inagaki and Katsuno, 2017). These waves play important roles in signal transduction (Xiong et al., 2016), cell division (Bement et al., 2015; Xiao et al., 2017) and cell migration (Gerisch et al., 2004; Weiner et al., 2007; Miao et al., 2017). Collective dynamics of cytoskeletal components not only amplify weaker signals originating from individual filament dynamics, but also serve as a readout for higher-order feedback in the system that may be challenging to assess solely at the level of individual filaments (Yang and Wu, 2018). Oscillatory behaviors offer temporal regularity, making them ideal tools for dissecting the crosstalk and feedback mechanisms of intertwined molecular networks (Kruse and Jülicher, 2005).

In this paper, we used oscillatory travelling waves on the cell cortex to examine the interplay between cytoskeletal networks. Our research focused on identifying key regulators important for the generation of actin waves. We confirmed the involvement of Cdc42 and FBP17 in mediating actin waves on the cortex of mast cells, consistent with previous findings (Wu et al., 2013, 2018; Xiong et al., 2016). Here we specifically explored the role of formins and microtubules as early regulators of cortical wave propagation. Our results revealed that the shrinkage of microtubules coincides with the release of FHDC1 from microtubules. This finding provided evidence for an antagonistic relationship between microtubule and actin polymerization in the context of wave dynamics. We further show the functional significance of these cytoskeletal waves in cellular processes. In the absence of FHDC1, wave propagation is perturbed, leading to defects in cell division, altered cell morphology and impaired locomotion. Collectively, our observations underscore the critical role of FHDC1 in mediating the crosstalk between microtubule and actin network in travelling waves.

## Results

### Formins are major nucleators of actin in cortical waves

In our previous studies, we have reported that resting and stimulated tumor mast cells (rat basophilic leukemia cells, or RBL cells) display cortical travelling waves of F-BAR protein FBP17 and actin (Wu et al., 2013). To gain insights into the propagation mechanisms of these actin waves, we captured the cortical dynamics of FBP17, and actin using total internal reflection fluorescence microscopy (TIRFM) at a fast acquisition rate (5 Hz) and showed that actin trailed FBP17 in travelling waves (Fig 1A, B, C, D). To visualize of the dynamics of individual actin puncta, we overexpressed mEos2-actin, a fluorescent protein that undergoes irreversible photo-conversion, shifting its emission peak from green (516 nm) to a red (581 nm) (Mckinney et al., 2009), and selectively illuminated a localized puncta in the mEos2-actin wave. We observed that the activated mEos2-actin remained in the same spot without any apparent spatial shift (Fig 1E), similar to FBP17 (Wu et al., 2018). This observation challenged the possibility of wave propagation by advection, where pre-existing actin molecules would simply move with the wave. Instead, our findings suggested that *de novo* nucleation of actin at the wave front likely plays a key role in supplying actin turnover during wave propagation. Notably, similar observations of *de novo* actin polymerization at the propagating wave front have also been reported in *Dictyostelium* cells (Bretschneider et al., 2009).

**Figure 1.**
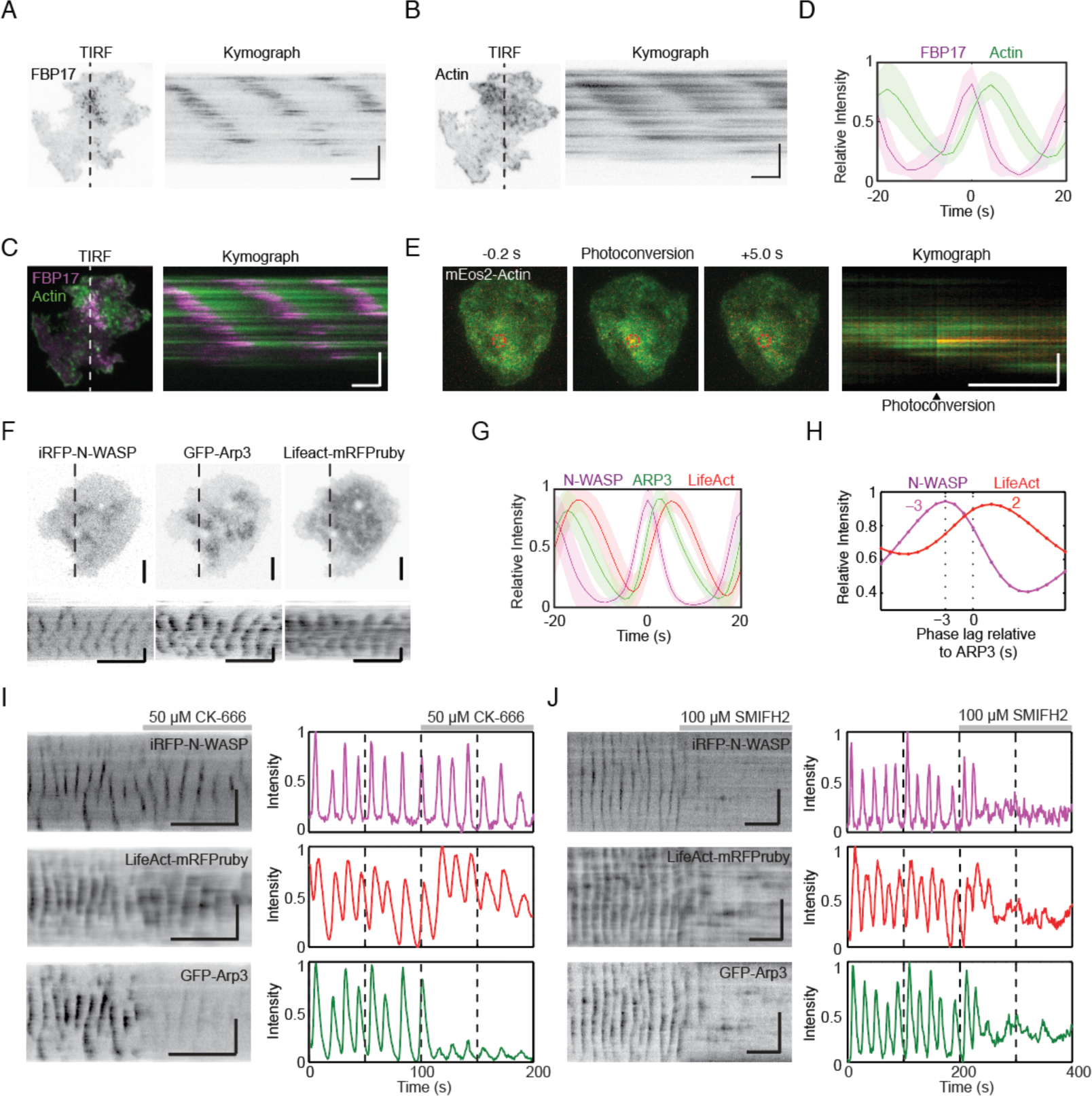
Actin nucleation in the waves is primarily mediated by formins. **(A, B)** Representative TIRFM micrographs and kymograph of a cell co-transfected with EGFP-FBP17 and actin-mCherry or LifeAct-mRFPruby (n=14; 3 independent experiments). Grayscale micrographs and kymographs are displayed with an inverted lookup table. Horizontal scale bars: 10 seconds. Vertical scale bars: 10 μm. **(C)** Two-color merge of TIRFM micrograph and kymograph of a representative cell co-transfected with EGFP-FBP17 (magenta) and actin-mCherry (green) from panels A and B. Horizontal scale bar: 10 seconds. Vertical scale bar: 10 μm. **(D)** Intensity profile of actin-mCherry (green) aligned with respect to EGFP-FBP17 (magenta). The solid lines represent the mean intensities, and the shaded region represent the standard deviations of the intensities. **(E)** Representative TIRFM micrographs and kymograph of cells transfected with mEos2-actin before and after photoconversion (n=98 cells; 12 independent experiments). The mEos2-actin punctum was photoconverted in the region indicated by the red circle. Horizontal scale bar: 10 seconds. Vertical scale bars: 10 μm. **(F)** Representative TIRM micrographs and kymographs of a cell co-transfected with iRFP-N-WASP, GFP-Arp3 and LifeAct-mRFPruby (n=74 cells; 7 independent experiments). Grayscale micrographs and kymographs are displayed with an inverted lookup table. Horizontal scale bars: 1 min. Vertical scale bars: 10 μm. **(G)** Representative intensity profile of GFP-Arp3 (green) and LifeAct-mRFPruby (red) aligned with respect to iRFP-N-WASP (n=74 cells; 7 independent experiments). The solid lines represent the mean intensities and the shaded regions represent the standard deviations of the intensities. **(H)** Representative cross correlation analysis of the time delay between the peak intensities of iRFP-N-WASP (magenta) and LifeAct-mRFPruby (red) to GFP-Arp3. **(I)** Representative kymographs and intensity profiles of a cell co-transfected with iRFP-N-WASP (magenta), LifeAct-mRFP-ruby (red), and GFP-Arp3 (green) and treated with CK-666 (n=8 cells; 2 independent experiments). Grayscale kymographs are shown with inverted lookup table. Horizontal scale bars: 1 min. Vertical scale bars: 10 μm. (J) Representative kymographs and intensity profiles of a cell co-transfected with iRFP-N-WASP (magenta), LifeAct-mRFP-ruby (red), and GFP-Arp3 (Green), and treated with SMIFH2 (n=23 cells; 6 independent experiments). Grayscale kymographs are shown with inverted lookup table. Horizontal scale bars: 1 min. Vertical scale bars: 10 μm.

To understand the nucleation mechanisms involved in actin dynamics during wave propagation, we first tested Arp2/3 complex, a major actin nucleation core. In our experiments, we overexpressed iRFP-N-WASP (an Arp2/3 activator), GFP-Arp3 (a component of the Arp2/3 complex) and LifeAct-mRFPruby (F-actin marker) and visualized their cortical dynamics. Our results show that N-WASP and Arp3 exhibit similar patterns of cortical waves as FBP17 and actin, respectively (Fig 1F). All presented as clusters of puncta within cortical waves. To gain insights into the temporal relationship between these proteins, we performed cross-correlation analysis on their intensity profile over time (Fig 1G). The analysis revealed that Arp3 puncta lagged the corresponding N-WASP puncta by approximately 3 s but preceded F-actin (labeled by LifeAct) by approximately 2 s (Fig 1H).

To determine the necessity of the Arp2/3 complex in the nucleation of actin waves, we used a chemical inhibitor of Arp2/3 CK-666. CK-666 keeps the Arp2/3 complex in an inactive state, preventing it from nucleating new actin filaments at the edges of existing filaments (Hetrick et al., 2013). Surprisingly, despite a significant decrease in the intensity and dynamics of Arp3 waves after the addition of CK-666 (Fig 1I), we observed that N-WASP and actin wave propagation persisted. In addition, there was a slight increase in actin intensity, indicating that Arp2/3 is not primarily responsible for the bulk of actin polymerization observed in actin waves (Fig 1I). These results motivated us to explore the involvement of formins in nucleating actin waves. We treated cells with a broad range formin inhibitor SMIFH2, which impedes the FH2 domains of formins and reduces their affinity for the barbed ends of actin filaments (Rizvi et al., 2009). Remarkably, both N-WASP and actin wave formation was completely abolished (Fig 1J). These pharmacological experiments suggest that formins are likely the major contributors to the nucleation Cdc42/FBP17-dependent actin waves in our system.

### FHDC1 is an early-phase regulator of cortical waves

To determine which formins are involved in actin wave formation, we examined the transcript expression levels of various formins in RBL cells using RNA-seq analysis (Fig 2A). Nine formins were expressed at varying levels, including Diaphanous Related Formin 1 (DIAPH1), Diaphanous Related Formin 2 (DIAPH2), Disheveled-Associated Activator of Morphogenesis 1 (DAAM1), Formin like 1 (FMNL1), Formin Homology 2 Domain Containing 1 (FHOD1), Formin like 2 (FMNL2), Formin 1 (FMN1), Formin 2 (FMN2) and FH2 Domain Containing 1 (FHDC1). We proceeded to overexpress each of these formins and visualize their cortical localizations. To eliminate any bias in the visual selection process, we performed Fast Fourier Transformations (FFTs) on the intensity profile, using the presence of distinct FFT peaks in the period of 0-40 s range as a criterion for local oscillations (waves). Based on this criterion, FMNL1, FHDC1 and DIAPH2 displayed oscillatory wave patterns in 60.0%, 41.8% and 35.7% of the cells with FBP17 waves, respectively. Other formins had wave-like patterns in only in a small subset of cells (Fig 2B). Interestingly, the percentage of cells exhibiting formin waves did not correlate with the individual transcript levels of the formins determined by RNA-seq analysis (Fig 2C). This suggests that the differences observed in wave participation reflect selective involvement in the pattern. Lastly, we also quantified the distribution of frequencies for the different formins and found no significant differences (Fig 2D).

**Figure 2.**
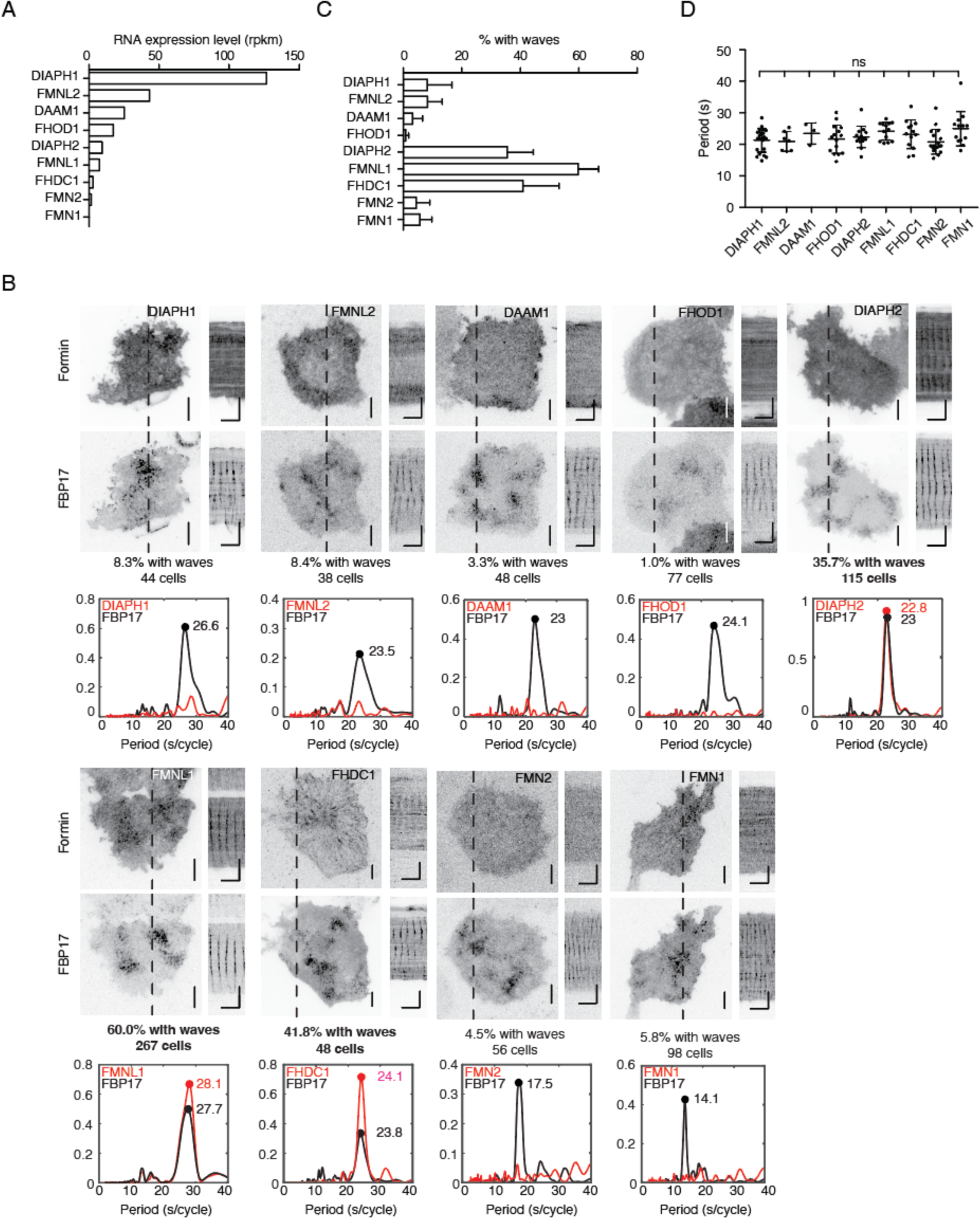
Localization of formins in the cortical travelling waves. **(A)** Formin transcripts expression levels in RBL cells determined by RNA-Seq analysis. Transcript expression levels were quantified as reads per kilobase per million mapped reads (RPKM). **(B)** Representative micrographs and kymographs of formin waves imaged over 2 minutes, along with FBP17 as wave marker. Grayscale micrographs and kymographs are displayed with an inverted lookup table. Horizontal scale bars: 1 minute. Vertical scale bars: 10 μm. Representative FFT plots of formin waves were generated. The three formins with the highest populations of cells displaying formin waves were highlighted in bold. **(C)** Quantification of the percentage of cells with formin travelling waves (4 – 23 independent experiments for each formin). **(D)** Quantification of frequency distributions of cortical formins’ oscillations in cells. Error bars represent the standard deviation.

Based on their regular involvement in the cortical waves in the majority of the cells, we further tested FMNL1, FHDC1 and DIAPH2 as key formins involved in actin wave formation. Notably, FHDC1 exhibited distinct localization patterns compared to FMNL1 and DIAPH2. FHDC1 was filamentous, while FMNL1 and DIAPH2 displayed a punctate pattern similar to that of FBP17 (Fig 2B). Beyond their localization patterns, FMNL1, FHDC1 and DIAPH2 waves also showed differences in their dynamics and phases. Co-expression of GFP-FMNL1 with mCherry-FBP17 showed strong co-localization, as FMNL1 shared a similar phase and temporal profile with FBP17 (Fig 3A – C). FMNL1 preceded actin in wave (Fig 3D). In contrast, imaging mCherry-FHDC1 and GFP-DIAPH2 with EGFP-FBP17 or mCherry-FBP17 suggested that FHDC1 and DIAPH2 peaked before FBP17 and exhibited an almost anti-phase relationship with actin (Fig 3E – L). Additionally, the assembly of FHDC1 and DIAPH2 on the membrane occurred at a slower rate compared to their disassembly, as indicated by gentler rising phases and sharper falling phases (Fig 3F, J). FHDC1 waves preceded FBP17 by approximately 4 s (−3.9 ± 0.2 s, 13 cells), while DIAPH2 waves preceded FBP17 by approximately 2 s (−1.8 ± 0.2 s, 14 cells). In contrast, FMNL1 peaks occurred on average approximately 1 s later than FBP17 (1.1 ± 0.1 s, 10 cells) (Fig 3M). Taken together, these data suggest that FMNL1 likely functions as an intermediator between N-WASP and actin, while FHDC1 and DIAPH2 act as early phase regulators of actin wave formation.

**Figure 3.**
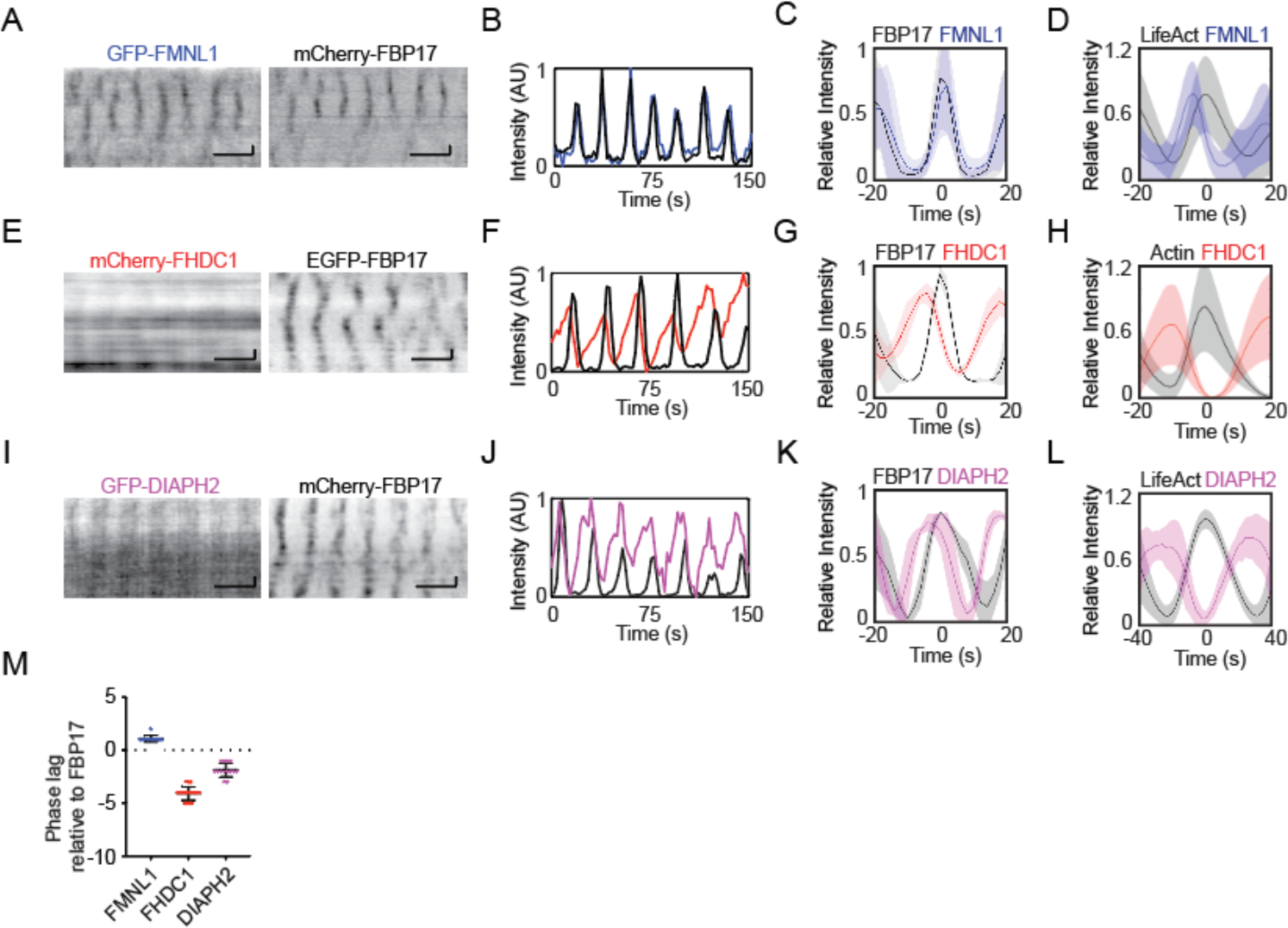
FHDC1 is an early phase regulator of cortical actin waves. **(A, B)** Representative kymographs and intensity profiles of a cell co-transfected with GFP-FMNL1 (blue) and mCherry-FBP17 (black) captured by TIRFM. Grayscale kymographs are shown with inverted lookup table. Horizontal scale bars: 1 minute. Vertical scale bars: 10 μm. **(C)** Representative intensity profile of GFP-FMNL1 (blue) aligned with respect to mCherry-FBP17 (black). The solid lines represent the mean intensities, and the shaded region represent the standard deviations of the intensities. **(D)** Representative intensity profile of GFP-FMNL1 (blue) aligned with respect to LifeAct-mRFP-ruby (black). The solid lines represent the mean intensities, and the shaded regions represent the standard deviations of the intensities. **(E, F)** Representative kymographs and intensity profile of a cell co-transfected with mCherry-FHDC1 (red) and EGFP-FBP17 (black) captured by TIRFM. Grayscale kymographs are shown with inverted lookup table. Horizontal scale bars: 1 min. Vertical scale bars: 10 μm. **(G)** Representative intensity profile of mCherry-FHDC1 (red) aligned with respect to EGFP-FBP17 (black). The solid lines represent the mean intensities, and the shaded regions represent the standard deviations of the intensities. **(H)** Representative intensity profile of mCherry-FHDC1 (red) aligned with respect to GFP-actin (black). The solid lines represent the mean intensities, and the shaded regions represent the standard deviations of the intensities. **(I, J)** Representative kymographs and intensity profiles of a cell co-transfected with GFP-DIAPH2 (magenta) and mCherry-FBP17 (black) captured by TIRFM. Grayscale kymographs are shown with inverted lookup table. Horizontal scale bars: 1 minute. Vertical scale bars: 10 μm. **(K)** Representative intensity profile of GFP-DIAPH2 (magenta) aligned with respect to EGFP-FBP17 (black). The solid lines represent the mean intensities, and the shaded regions represent the standard deviations of the intensities. **(L)** Representative intensity profile of GFP-DIAPH2 (magenta) aligned with respect to LifeAct-mRFP-ruby (black). The solid lines represent the mean intensities, and the shaded regions represent the standard deviations of the intensities. **(M)** Quantification of the phase lag of FMNL1, FHDC1 and DIAPH2 with respect to FBP17.

### Interactions of FHDC1 with both actin and microtubule regulate wave propagation

We focused on investigating FHDC1 in our study due to its early recruitment to cortical waves and distinct filamentous localization pattern, which distinguishes it from other formins. Unlike typical formins that have conserved formin homology (FH) domains at their C-terminal, FHDC1 exhibits a unique arrangement. Its FH1 and FH2 domains are in the N-terminal half, while the C-terminal half contains a microtubule-binding domain (FHDC1 958-1143) (Young et al., 2008). Interestingly, this C-terminal microtubule-binding domain is predicted to be intrinsically disordered, suggesting a flexible and dynamic nature of this region (Fig 4A). This unique arrangement and the intrinsic disorder prediction raise intriguing questions about the functional properties and regulatory mechanisms of FHDC1 in actin wave propagation.

**Figure 4.**
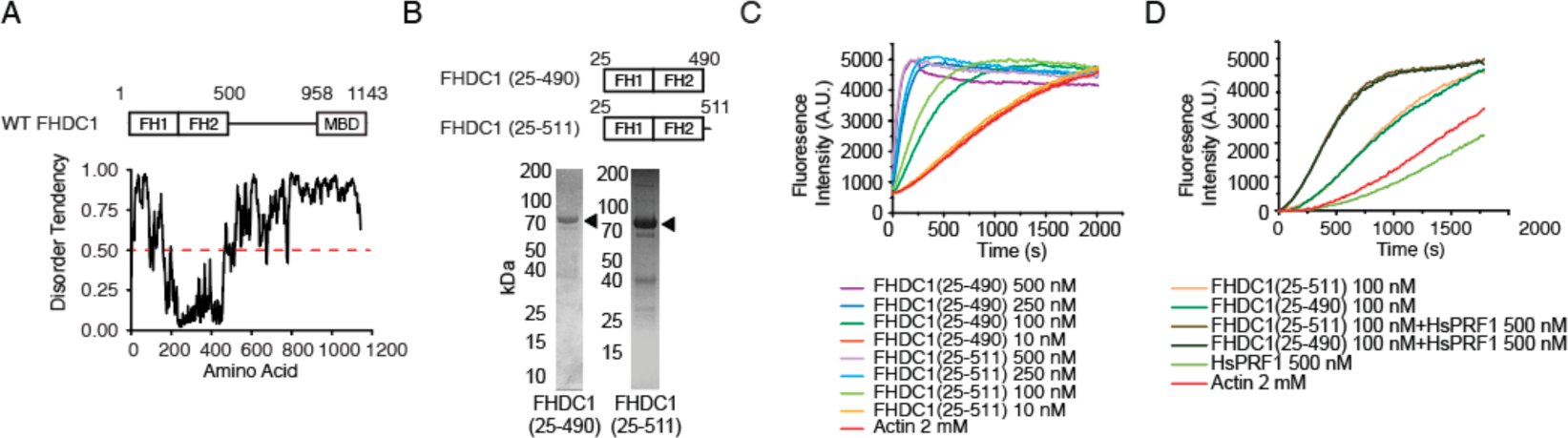
The FH domains of FHDC1 are sufficient for actin polymerization. **(A)** Diagram illustrating the prediction of intrinsically disorder region (IDR) in the full-length FHDC1 using the IUPRED2 algorithm (https://iupred2a.elte.hu/). Regions with an IDR score above 0.5 are considered to be intrinsically disordered. **(B)** Domain organization of truncated mutant constructs of FHDC1 used for Pyrene-actin polymerization assay. Purified FHDC1 truncating variants analyzed using Coomassie blue-stained SDS-PAGE gels. **(C)** Pyrene-actin polymerization assay performed using 2 μM actin (5% pyrene-labeled) in the presence of the indicated concentration of FHDC1(25-490) and FHDC1(25-511) truncated variants. **(D)** Pyrene-actin polymerization assay using 2 μM actin (5% pyrene-labeled) in the presence of the 100 nM FHDC1(25-490), 100 nM FHDC1(25-511) and 500 nM HsPRF1.

To further explore the functional properties of FHDC1, including its ability to polymerize actin, we purified two truncation mutants of FHDC1, namely FHDC1 (25-490) and FHDC1 (25-511) (Fig. 4B), and performed *in vitro* pyrene actin polymerization assays. The results confirmed that both proteins were capable of nucleating actin polymerization (Fig 4C). Moreover, the nucleation activity could be accelerated by Profilin-1 (HsPRF1) (Fig 4D), a known regulator of actin dynamics.

Next, to determine the specific role of FHDC1 in mediating wave propagation, we performed a knockdown targeting FHDC1. The knockdown resulted in a significant reduction in the percentage of cells displaying waves as marked by EGFP-CBD (Cdc42-binding-domain), from 84.8 ± 4.6% to 32.6 ± 4.6% (Scrambled shRNA: n=4, 183 cells; FHDC1 knockdown: n=9, 273 cells, P<0.0001, one-way ANOVA; adjusted P<0.0001, Sidak’s multiple comparison post hoc test) (Fig 5A, B), suggesting that FHDC1 plays an important role in facilitating wave propagation. To further characterize the relationship between FHDC1 and microtubules, we generated truncated constructs encompassing different regions of FHDC1 (Fig 5C) and examined the co-localization and wave dynamics of α-tubulin and FHDC1. Interestingly, we observed that the full-length FHDC1 not only co-localized with α-tubulin but also displayed similar wave dynamics (Fig 5D), indicating that FHDC1 is associated with microtubules and is involved in wave propagation. Unsurprisingly, the FHDC1 truncated mutant lacking FH domains (FHDC1 501-1143) also localized to microtubules (Fig 5E). However, the central unstructured region (FHDC1 501-958), which has not been previously characterized, was found to be sufficient for microtubule association (Fig 5F). These findings confirmed the association of microtubule with FHDC1 and suggest that the region responsible for FHDC1’s binding to microtubule may extend beyond the previously identified microtubule-binding domain.

**Figure 5.**
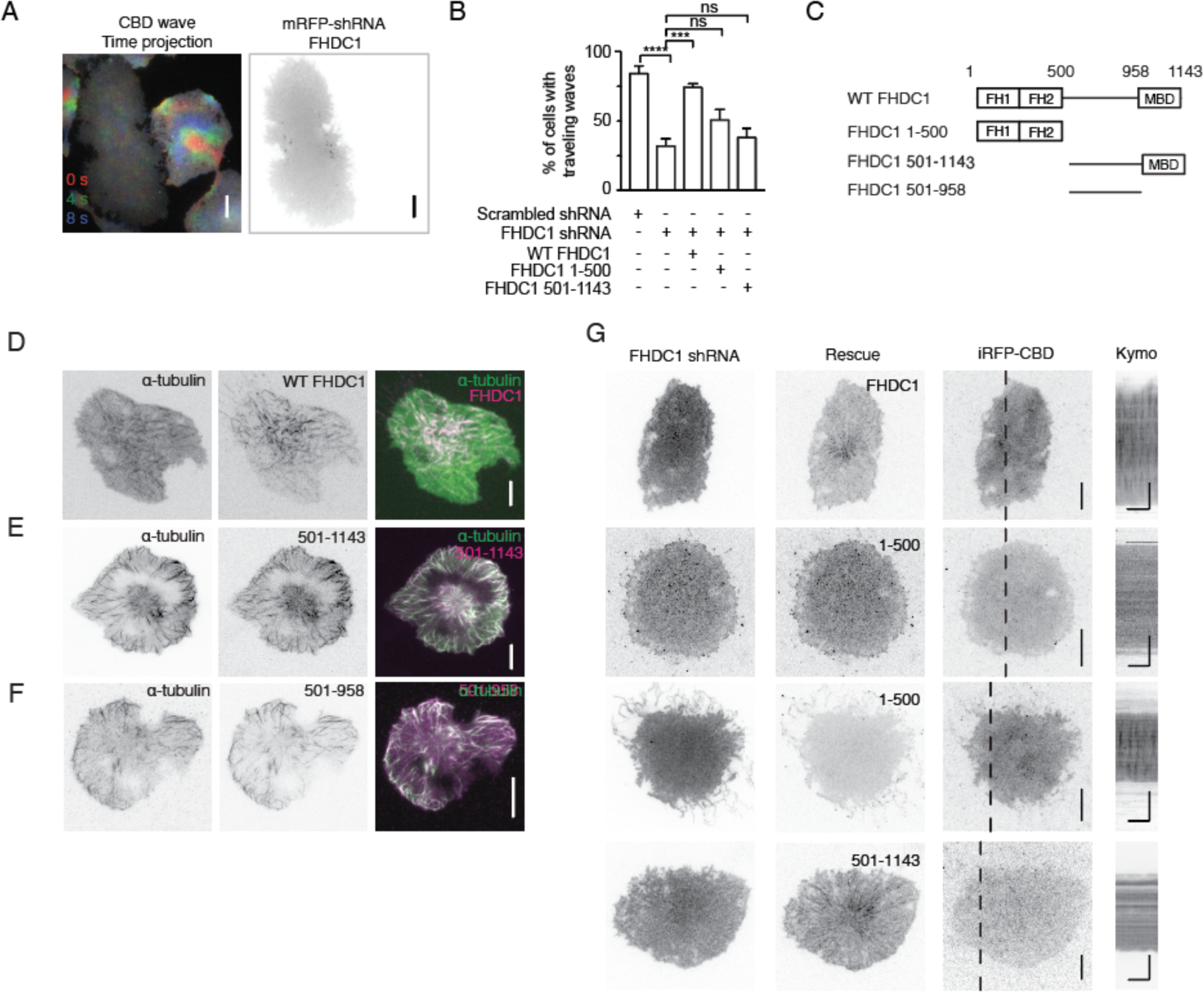
Both actin and microtubule interacting domains of FHDC1 are required for wave propagation. **(A)** Left: Representative time projection image of CBD wave propagation over 8 seconds (n=242; 8 independent experiments). The pseudocolors representing each frame are as follows: Red: 0 seconds, Green: + 4 seconds. Blue: + 8 seconds. Right: Co-expression of FHDC1 shRNA constructs in an EGFP-CBD stable cell line abolished propagation of CBD wave. Grayscale micrographs are shown with inverted lookup table. Scale bar: 10 μm. **(B)** Quantification of the percentage of cells with CBD travelling waves following transfection with FHDC1 shRNA (n=242; 8 independent experiments) or scrambled shRNA (n=183; 4 independent experiments), compared with rescue by full length GFP-FHDC1 (n=108; 3 independent experiments), GFP-FHDC1 (1-500) (n=92; 3 independent experiments), or GFP-FHDC1 (501-1143) (n=86; 3 independent experiments). **(C)** Domain organization of wild type and truncated mutant constructs of FHDC1 used. **(D)** Representative micrograph of a cell co-transfected with GFP-α-tubulin and mCherry-FHDC1 captured by TIRFM. Grayscale micrographs and are shown with inverted lookup table. Scale bar: 10 μm. The merged micrograph is represented by the following pseudocolors: GFP-α-tubulin (green), mCherry-FHDC1 (magenta). **(E)** Representative micrograph of a cell co-transfected with mCherry-α-tubulin and GFP-FHDC1 (501-1143) captured by TIRFM. Grayscale micrographs and are shown with inverted lookup table. Scale bar: 10 μm. The merged micrograph is represented by the following pseudocolors: mCherry-α-tubulin (green), GFP-FHDC1 (501-1143) (magenta). **(F)** Representative micrograph of a cell co-transfected with mCherry-α-tubulin and GFP-FHDC1 (501-958) captured by TIRFM. Grayscale micrographs and are shown with inverted lookup table. Scale bar: 10 μm. The merged micrograph is represented by the following pseudocolors: mCherry-α-tubulin (green), GFP-FHDC1 (501-958) (magenta). **(G)** Representative micrographs and kymographs of cells co-transfected with pRFP-C-RS-FHDC1 shRNA constructs, iRFP-CBD and either full length GFP-FHDC1, GFP-FHDC1 (1-500) or GFP-FHDC1 (501-1143). The percentage shown represent cells with CBD waves observed. Grayscale micrographs and kymographs are shown with inverted lookup table. Horizontal scale bars: 10 seconds. Vertical scale bars: 10 μm.

To validate the importance of FHDC1 in wave propagation, we attempted to rescue the effects of FHDC1 knockdown to wave propagation by re-introducing an shRNA resistant FHDC1 construct. The re-introduction of FHDC1 successfully restored wave propagation to levels observed prior to the knockdown (74.8 ± 2.1% cells, n=3, 108 cells, P<0.0001, one-way ANOVA; adjusted P=0.0002, Sidak’s multiple comparison post hoc test), confirming the importance of FHDC1 in facilitating wave propagation. In contrast, re-introducing FHDC1 1-500 alone failed to restore wave propagation in approximately 50% of the knockdown cells imaged (51.3 ± 7.0% cells, n=3, 92 cells, P<0.0001, one-way ANOVA; adjusted P=0.1271, Sidak’s multiple comparison post hoc test) (Fig 5B, G). We ruled out the possibility that the lack of rescue was due to the choice of the FH domain as both FHDC1 (25-490) and FHDC1 (25-511) showed competence in nucleating actin in the pyrene assay. Hence, the primary reason for the failed rescue can be attributed to the absence of microtubule binding. Similarly, the microtubule-binding segments of FHDC1 (FHDC1 501-1143) also failed to rescue wave propagation in knockdown cells (38.7 ± 5.9% cells, n=3, 86 cells, P<0.0001, one-way ANOVA; adjusted P=0.9448, Sidak’s multiple comparison post hoc test) (Fig 5B, G), indicating actin nucleation was also essential for wave propagation. These data collectively indicate that both the FH domains and microtubule-binding domains of FHDC1 are important in mediating wave propagation.

### Microtubule depolymerization regulates actin waves

We proceeded to investigate how the dynamics tubulin waves could be linked to actin waves. Co-imaging GFP-α-tubulin with iRFP-N-WASP revealed that the disappearance of α-tubulin coincided with the appearance of N-WASP waves (Fig 6A, B). Similarly, when LifeAct-mRubyRFP was co-imaged with GFP-α-tubulin, actin waves were observed to trail behind α-tubulin waves (Fig 6C, D). While there was some degree of overlap between the signals of N-WASP and microtubules (Fig 6B), the peaks of actin were found to be anti-phased with microtubule waves (Fig 6D). To confirm that disappearance of the microtubule was indeed due to depolymerization rather than leaving the TIRF field, we tracked the dynamics of microtubules plus ends by co-expressing GFP-EB1 with mCherry-α-tubulin and iRFP-N-WASP. The arrival of N-WASP coincided with the shrinkage of microtubule. and microtubule underwent new cycles of polymerization after each wave of N-WASP passed (Fig 6E). Next, we aimed to determine the functional importance of microtubule dynamics. Preventing microtubule polymerization using nocodazole did not affect wave dynamics (Fig 7A). However, stabilizing microtubules with taxol resulted in an instant inhibition of wave propagation (Fig 7B). Interestingly, in cells with taxol-stabilized microtubules, the recovery of mCherry-FHDC1 fluorescence after photo-bleaching was significantly reduced compared to untreated cells (Fig 7C, D), indicating slowed release of FHDC1. Taken together, we propose that depolymerization of microtubule can be utilized to regulate cortical waves through the release of FHDC1 (Fig 7E).

**Figure 6.**
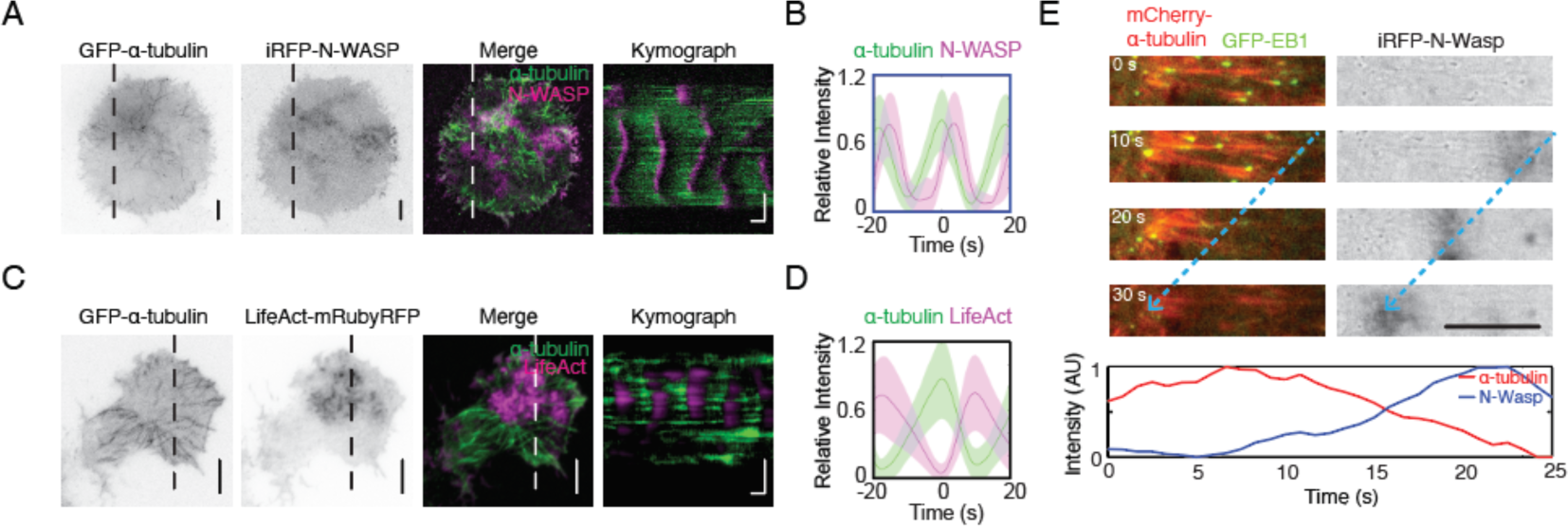
Microtubule depolymerisation coincides with actin waves. **(A)** Representative TIRFM micrographs and kymograph of a cell co-transfected with iRFP-N-WASP and GFP-α-tubulin (30 cells). Grayscale micrographs are shown with inverted lookup table. Horizontal scale bar: 10 seconds. Vertical scale bars: 10 μm. The merged micrograph and kymograph are represented by the following pseudocolors: GFP-α-tubulin (green), iRFP-N-WASP (magenta). **(B)** Representative intensity profile of iRFP-N-WASP (magenta) aligned with respect to GFP-α-tubulin (green). The solid lines represent the mean intensities, and the shaded regions represent the standard deviations of the intensities. **(C)** Representative TIRFM micrographs and kymograph of a cell co-transfected with GFP-α-tubulin and LifeAct-mRubyRFP (12 cells). Grayscale micrographs are shown with inverted lookup table. Horizontal scale bar: 10 seconds. Vertical scale bars: 10 μm. The merged micrograph and kymograph are represented by the following pseudocolors: GFP-α-tubulin (green), LifeAct-mRubyRFP (magenta). **(D)** Representative intensity profile of LifeAct-mRubyRFP (magenta) aligned with respect to GFP-α-tubulin (green). The solid lines represent the mean intensities, and the shaded regions represent the standard deviations of the intensities. **(E)** Top: Representative merged montages of microtubule shrinkage are displayed with the following pseudocolors: GFP-EB1 (green) and mCherry-α-tubulin (red). Grayscale montage of iRFP-N-WASP are shown with inverted lookup table. Scale bar: 10 μm. Blue dotted arrows indicate the direction of iRFP-N-WASP wave propagation and mCherry-α-tubulin depolymerization. Bottom: Intensity profile of iRFP-N-WASP (blue) and mCherry-α-tubulin (red) is shown for the montage captured over 25 seconds.

**Figure 7.**
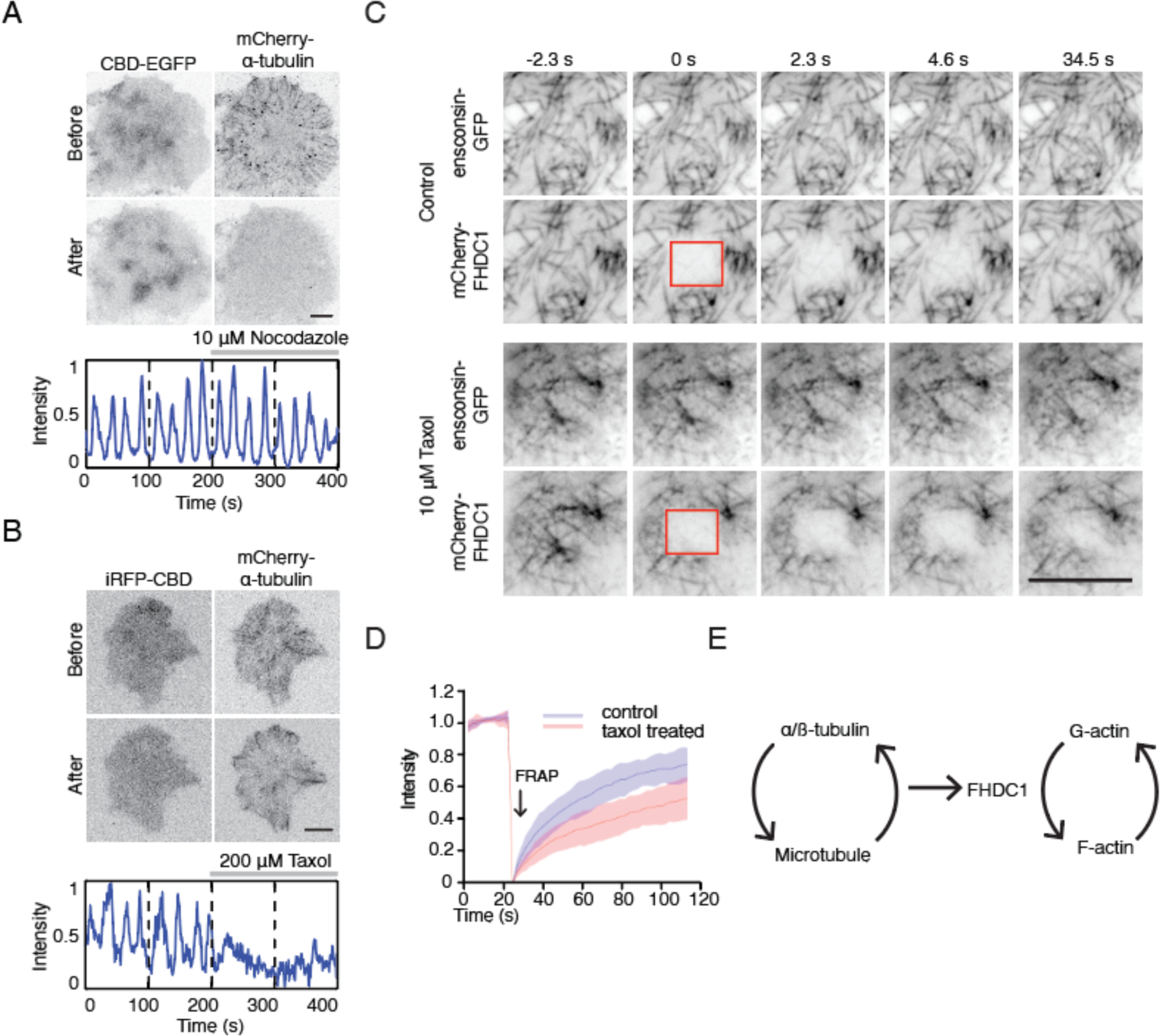
Depolymerization of microtubule regulate cortical waves through FHDC1. **(A)** Top: Representative micrographs of cells stably expressing CBD-EGFP co-transfected with mCherry-α-tubulin before and after 10 μM nocodazole treatment (n=13; 5 independent experiments). Bottom: Intensity profile of CBD-EGFP of the above cell before and after nocodazole treatment over 400 seconds. Nocodazole treatment is indicated by the gray bar above the intensity profile. **(B)** Top: Representative micrographs of cells co-transfected with iRFP-CBD and GFP-α-tubulin before and after 200 μM taxol treatment (n=5, 3 independent experiments). Bottom: Intensity profile of CBD-EGFP of the above cell before and after taxol treatment over 400 seconds. Taxol treatment is indicated by the gray bar above the intensity profile. **(C)** Representative micrographs of mCherry-FHDC1 with ensconsin-GFP (microtubule marker) in 10 μM taxol medium or control medium. The photobleached area by 561nm laser is shown by red boxes. Scale bar: 10 μm. **(D)** Intensity profile of fluorescence recovery after photobleaching (FRAP) of mCherry-FHDC1 in 8 cells for each treatment. Shadowed area denotes Standard Deviation. **(E)** Model of crosstalk between microtubule and actin regulation by FHDC1.

### FHDC1 knockdown cells display defects in cell polarity, locomotion and division

To assess the physiological impact of perturbing wave dynamics by FHDC1 knockdown, we next characterized the cellular defects associated with FHDC1 depletion. One significant phenotype observed in FHDC1 knockdown cells was an increase in the number of cell protrusions (Fig 8A). Wild type cells generally exhibited two cell protrusions on average (2.2 ± 0.1 protrusions, 51 cells). However, upon FHDC1 knockdown, the number of observed protrusions significantly increased, with some cells displaying up to eight protrusions observed (3.8 ± 0.2 protrusions, 51 cells) (Fig 8B). This elevation suggests a reduced ability of FHDC1 knockdown cells to establish cell polarity. Furthermore, cell locomotion was impaired as indicated by a decrease in cell velocity from 14.7 ± 0.6 μm/min in wild type cells to 10.5 ± 0.6 μm/min in FHDC1 knockdown cells (P<0.0001, student’s t-test) (Fig 8C). FHDC1 knockdown also resulted in a significant reduction in the number of successful cell division events. In wild type cells, 80.1% ± 2.9% of the cells successful underwent division within a 24 h period (n=3, 194 cells). Cells transfected with scrambled shRNA exhibited a comparable rate of successful division (83.7% ± 8.7%, n=2, 93 cells). In contrast, cells transfected with FHDC1 shRNA demonstrated a substantial decrease, with only 25.5% ± 3.5% of the cells undergoing successful division (n=3, 149 cells) (P=0.0051, student’s t-test) (Fig 8D). The results suggest that FHDC1 may play a role in coordinating the actin and microtubule cytoskeletons during cell division.

**Figure 8.**
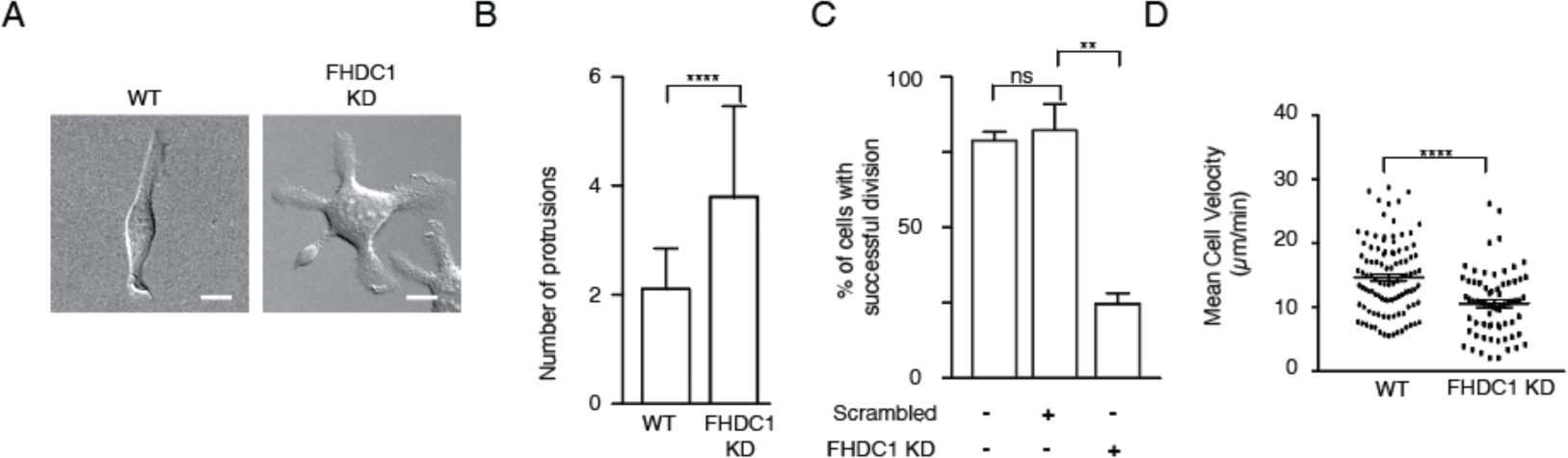
FHDC1 knockdown changes cell motility and impairs cell division. **(A)** FHDC1 knockdown cells present increased numbers of cell protrusions. Representative micrographs of wild type cell and cell transfected with FHDC1 shRNA imaged by DIC. Scale bar: 10 μm. **(B)** Quantification for number of cell protrusions between wild type cells and FHDC1 knockdown cells. **(C)** Quantification of cell velocity between wild type cells and FHDC1 knockdown cells. **(D)** Quantification of percentage of cells with successful cell division between wild type cells and FHDC1 knockdown cells.

## Discussion

In this paper, we have demonstrated that formins play a crucial role in regulating actin wave propagation. Recent studies have also highlighted the importance of formins in this process (Jasnin et al., 2016; Ecke et al., 2020). In addition, we made an observation of travelling waves of the formin FHDC1 and microtubules on the plasma membrane, which have not been previously observed. These observations have allowed us to identify microtubules and FHDC1 as key regulators of actin waves and offer a unique opportunity to dissect the crosstalk between these cytoskeletal networks in living cells.

The crosstalk of actin and microtubule dynamics is regulated through various mechanisms (Dogterom and Koenderink, 2018). One mode of regulation involves direct physical crosslinking of actin and microtubule. Protein such as tau and spectraplakins are examples that possess both actin and microtubule-binding domains that allow them to directly link and align actin filaments with microtubules (Wu et al., 2008; Fontela et al., 2017). This physical interaction promotes co-alignment and coordination between the two cytoskeletal systems. Another form of crosstalk involves competition between actin and microtubules for binding to common interacting proteins (Henty-Ridilla et al., 2016). There are proteins that may have the ability to interact with both actin and microtubules and act as shared regulators. The binding of these proteins to either actin or microtubules can influence the dynamics of the respective cytoskeletal network, thereby indirectly affecting the dynamics of the other network. Additionally, actin and microtubules can share a common upstream regulator that coordinate their dynamics. RhoGTPases, such as Rho, Cdc42, and Rac are key regulators of both actin polymerization and microtubule dynamics. They can activate signaling pathways that mediate actin stress fiber formation or facilitate microtubule stabilization and organization (Waterman-storer et al., 1999; Fukata et al., 2002; Krendel et al., 2002; Graessl et al., 2017).

There is growing evidence indicating that formins are prominent regulators of both actin and microtubules (Chesarone and Goode, 2010; Bartolini and Gunderson, 2011). Formins can interact with microtubules through various mechanisms. Some formins, such as DIAPH1, DIAPH2 and DIAPH3, bind to microtubule using their FH domains, which are primarily involved in actin polymerization (Palazzo et al., 2001; Bartolini et al., 2005; Gaillard et al., 2011; Daou et al., 2014). Other formins, such as FMN1 and FHDC1, have separate microtubule-binding domains that allow them to directly interact with microtubules (Zhou et al., 2006; Young et al., 2008). Additionally, formins like DAAM1 may use a combination of FH domains and microtubule-binding domains to interact with both actin and microtubules (Szikora et al., 2017). Formins can bind to microtubules at different sites, such as along the entire microtubules (FHDC1, FMN2) (Young et al., 2008; Kwon et al., 2011) or accumulate as puncta along the sides or ends of microtubules (DIAPH1) (Henty-Ridilla et al., 2016). Furthermore, formins have been shown to modulate post-translational modifications of microtubules, such as detyrosination (Andrés-Delgado et al., 2012) and acetylation (Thurston et al., 2012; Fernández and Miguel, 2018). However, the functional implications of these mechanistic variances in formin-microtubule interactions are not yet fully understood.

In our study, we found that all nine formins expressed in RBL cells could have the potential to be involved in actin waves, although some may only participate in a small subset of cells with actin waves. This highlights the complexity of cortical dynamics, but the importance of formins in actin wave regulation is likely general. In *Dictyostelium* cells, inhibiting formins has been shown to suppress localized actin accumulation on perforations that are initiated by wave propagation (Jasnin et al., 2016). Among the nine formins, six (DAAM1, DIAPH1, DIAPH2, FHDC1, FMN1 and FMN2) have the potential to interact with microtubule. We focused on FHDC1 due to its robust involvement in actin waves and co-localization with microtubules, which distinguishes it from the other formins. FHDC1 exhibits a significant phase shift ahead of actin, suggesting that its actin nucleation function is likely inhibited when associated with microtubules. This unique characteristic makes FHDC1 an ideal localized source of active nucleator that can be released with precise temporal control. While many studies have demonstrated cooperative roles of actin and microtubule coordination in cell division and migration, our proposed antagonistic role between actin and microtubules is distinct (Waterman-storer et al., 1999; Wu et al., 2008, 2018; Bement et al., 2015; Ning et al., 2016; Winans et al., 2016). We found that cortical regions with stable microtubules lack actin waves, and stable microtubules inhibit actin wave formation. We hypothesize that this antagonistic relationship is mediated through the release of FHDC1. The theoretical plausibility of such antagonistic relationships is supported by the fact that microtubule binding or stability can inhibit the actin nucleation activity of some formins including DIAPH1 and DIAPH3 (Gaillard et al., 2011; Bartolini et al., 2012). Similar antagonism has been observed in other lymphocytes (Inoue et al., 2018).

Our findings reveal that while microtubule dynamics can regulate actin waves, microtubules are not essential for actin waves propagation. In the absence of microtubules upon complete depolymerization, Cdc42-dependent actin waves can still propagate. Interestingly, we have observed that the depolymerization of microtubule enhances RhoA-dependent actin wave formation, as previously reported by us and other research groups (Krendel et al., 2002; Graessl et al., 2017; Xiao et al., 2017; Tong and Wu, 2023). Based on these observations, we propose three distinct regimes of actin/microtubule waves: (I) stable microtubule without actin waves, (II) dynamic microtubule waves accompanied by Cdc42-dependent actin waves, and (III) no microtubule with both Cdc42 and RhoA-dependent actin waves. These results highlight the remarkable plasticity displayed by cells in coordinating its cytoskeletal networks. Depending on the compositions of regulatory proteins involved, the cortex structure can be fine-tuned and dynamically remodeled to adapt different cellular requirements.

## Materials and Methods

### Cell Culture, transfection and drug treatments

Rat Basophilic Leukemia (RBL-2H3) cells were cultured as monolayer in MEM growth medium (Life Technologies, Carlsbad, CA) supplemented with 20% heat-inactivated Fetal Bovine Serum (Sigma-Aldrich, St Louis, MO). The cells were harvested with TrypLE Express (Life Technologies, Carlsbad, CA) three days after passage. Transient transfections were performed on cells in suspension by subjecting them to two pulses at 1200 mV for 20 ms using Neon Transfection Electroporator (Life Technologies, Carlsbad, CA). After transfection, cells were plated in 35 mm glass-bottom Petri dishes (In vitro scientific and MatTek) at sub-confluent densities and sensitized with anti-DNP IgE (Life Technologies, Carlsbad, CA) at 0.5 ug/mL. The cells were maintained overnight in a humidified incubator at 37°C. For experiments involving the use of inhibitors, the inhibitors were diluted from stock solutions and added to the cells at the following final concentrations: CK-666 (50 μM, Sigma-Aldrich, St Louis, MO), SMIFH2 (100 μM, Sigma-Aldrich, St Louis, MO), Taxol (100 μM, Sigma-Aldrich, St Louis, MO), Nocodazole (10 μM, Sigma-Aldrich, St Louis, MO).

To achieve targeted FHDC1 knockdowns, a combination of four unique 29mer pRFP-C-RS-FHDC1 short hairpin RNA (shRNA) constructs (TF705017A, B, C, D, OriGene Technologies Inc, Rockville, MD) was introduced into the cells. For controls, scrambled shRNA (TR30012, OriGene Technologies Inc, Rockville, MD) was used instead. Prior to imaging, the cells were incubated at 37°C in a humidified CO_2_ incubator for 48 hours to allow for gene silencing and ensure maximum clearance of endogenous FHDC1. The selection of FHDC1 knockdown cells was based on the expression of turboRFP. To rescue the FHDC1 knockdown phenotype, shRNA-resistant constructs carrying human FHDC1 sequences (GFP-FHDC1, GFP-FHDC1(1-500) and GFP-FHDC1(501-1143) were co-transfected with the FHDC1 shRNAs. Additionally, iRFP-CBD was used as a readout for waves. After transfection, cells were incubated at 37°C in a humidified CO_2_ incubator for 48 hours to allow for gene silencing and the expression of human FHDC1 before imaging.

### Molecular Cloning and plasmids

To generate iRFP-N-WASP, the iRFP fluorescent tag was subcloned into the pEGFP-C1 backbone vector, replacing the EGFP, using kpnI site. The N-WASP sequence was then subcloned from GFP-N-WASP into the modified pEGFP-C1 vector. For construction of mCherry-FBP17, the FBP17 was subcloned using XhoI and EcoRI sites from EGFP-FBP17 into pmCherry-C1 backbone vector. To create FBP17-mEos3, the FBP17 sequence was subcloned from EGFP-FBP17 into mEos3.2-N1 vector using EcoRI and BamHI restriction sites.. Truncated GFP-FHDC1 1-500 and GFP-FHDC1 501-1143 mutant were generated by subcloning the respective sequences into pEGFP-C1 vector using the KpnI and BamHI restriction sites. The source of these constructs was the mCherry-FHDC1. Constructs for the following proteins were obtained as kind gifts: LifeAct-mRuby from Dr Roland Wedlich-Soldner (Max Planck Institute of Biochemistry, Martinsried, Germany); mEos2-Actin-7 from Dr. Michael W. Davidson (Florida State University, Tallahassee, FL); GFP-α-tubulin from Dr Yih-Cherng Liou, (National University of Singapore, Singapore); GFP-Arp3 from Dr Dorothy Schafer (University of Massachusetts Medical School, Worcester, MA); FBP17-EGFP and mCherry-actin from Dr Pietro De Camilli; GFP-DIAPH1, GFP-DIAPH2, FMNL2-GFP, DAMM1-GFP and FHOD1-GFP from Dr Alexander Bershadsky (Mechanobiology Institute, Singapore); FMN1-GFP from Dr Michael Sheetz (Mechanobiology Institute, Singapore); GFP-FMN2 from Dr Sonia Rocha (University of Dundee, Dundee, Scotland, UK); FMNL1-GFP from Dr Michael Rosen (UT Southwestern Medical Center, Dallas, TX); mCherry-FHDC1 from Dr John Copeland (University of Ottawa, Ottawa, Ontario, Canada).

### Microscopy

Before imaging, the medium in the dishes was replaced with Tyrodes’s imaging buffer (20mM HEPES (pH 7.4), 135mM NaCl, 5.0mM KCl, 1.8mM CaCl2, 1.0mM MgCl2 and 5.6mM glucose). The cells were then transferred to a heated microscope stage (Live Cell Instrument, Seoul, South Korea) maintained at 37 °C throughout the experiments. For TIRFM imaging of live cell cortical dynamics, a Nikon Ti-E inverted microscope (Nikon, Shinagawa, Tokyo) equipped with a perfect focus system to prevent focus drift was used. The microscope is also equipped with an iLAS2 motorized TIRF illuminator (Roper Scientific, Evry Cedex, France) and either an Evolve 512 EMCCD camera (Photometrics, Tucson, AZ) (16 bit, pixel size 16 μm) or Prime95b sCMOS camera (Photometrics, Tucson, AZ) (16 bit, pixel size 11 μm). Objective lenses from Nikon’s CFI Apochromat TIRF Series (100xH N.A. 1.49 Oil; 60xH N.A. 1.49 Oil) were used for image acuisition. Multi-channel imaging of samples was achieved by the sequential exciting the samples with 491 nm (100 mW), 561 nm (100 mW) and 642 nm (100 mW) lasers, reflected from a quad-bandpass dichroic mirror (Di01-R405/488/561/635, Semrock, Rochester, NY) located on a Ludl emission filter wheel (Carl Zeiss AG, Oberkochen, Germany). The microscope was controlled using MetaMorph software (Version 7.8.6.0) (Molecular Devices, LLC, Suunyvale, CA). During image acquisition, the samples were maintained at 37 °C using an on-stage incubator system (Live Cell Instrument, Seoul, South Korea). Prior to or during imaging, stimulation was performed by adding 80 ng/mL of DNP-BSA, a multivalent antigen that stimulates an antigen response. To ensure cell viability during long term imaging lasting more than one hour, the spent media was replaced with fresh media. Additionally, 5% humidified CO_2_ was maintained. For photo-conversion of mEos2-Actin, the 405 nm laser was set to its maximum power and pulsed on a single punctum for 20 ms. The photo-conversion process was controlled using the ‘On Fly’ module of the iLAS2 controller. To perform Fluorescence Recovery After Photobleaching (FRAP) on mCherry-FHDC1 and ensconsin-GFP treated with taxol and nocodazole, a region of interest (ROI) measuring 7 x 6 μm was selected, and photobleaching was carried out using the 561 nm laser set to maximum power. The FRAP experiments were controlled using the ‘On-Fly’ module of the iLAS2 controller.

To study the functional effects of FHDC1 knockdown on cell division and locomotion, wide-field epi-fluorescence and DIC microscopy were used in tandem. Images acquisition was performed using a Nikon Ti-E inverted microscope (Shinagawa, Tokyo, Japan). The microscope was equipped with an X-Cite 120LED microscope light source (Excelitas Technologies Corp, Waltham, MA, 370–700 nm) and an ORCA-Flash 4.0 V2 Digital CMOS camera C11440-22CU (Hamamatsu, 16 bit, pixel size 6.5 μm). All images were acquired using an objective len from Nikon’s CFI Plan Apochromat Lambda (λ) Series (40x N.A. 0.95). The microscope was controlled using the NIS-Elements AR 4.20 software (Nikon, Shinagawa, Tokyo). To ensure cell survival during image acquisition, the samples were maintained at 37 °C with a 5% humidified CO2 environment.

### Protein Purification

Truncating variants of FHDC1 were fused with an N-terminal GST-His6 tag in pNIC-GST vector using ligation-independent cloning (LIC) technology. HsPRF1 was fused with N-terminal 8xHis tag in a modified version of pET-21d (+) pSY5 vector. The respective bacterial expression vectors were transformed into E.coli BL21 (DE3) Rosetta T1R and cultured in Terrific Broth supplemented with 8 g/L glycerol. The cultures were incubated at 37°C with shaking at 200 rpm overnight and induced with 0.5 mM IPTG for protein expression. The induced cultures were further incubated overnight at 18 °C. Cells were harvested and resuspended in lysis buffer (100 mM HEPES, 500 mM NaCl, 10 mM Imidazole, 10 % glycerol, 0.5 mM TCEP, pH 8.0) supplemented with protease inhibitor cocktail set III, EDTA free (diluted 1000x in lysis buffer, Calbiochem, USA), and benzonase (Merck, USA) at a final concentration of 5 μL per liter of culture. The re-suspended cell pellet was sonicated using a Sonics Vibra-cell at 70% amplitude for 3 minutes on ice with a cycle of 3 seconds on/off. The lysate was clarified by centrifugation at 47,000 g, 4°C for 25 minutes. The supernatants were filtered through 1.2 μm syringe filters and loaded onto an AKTA Xpress system (GE Healthcare). The lysates were loaded on IMAC columns and. washed with binding buffer (20 mM HEPES, 500 mM NaCl, 25 mM Imidazole, 10 % (v/v) glycerol, 0.5 mM TCEP, pH 7.5). Gradient elution was performed using elution buffer (20 mM HEPES, 500 mM NaCl, 500 mM Imidazole, 10 % (v/v) glycerol, 0.5 mM TCEP, pH 7.5). The eluted proteins were collected, stored in sample loops on the system and injected into a Gel Filtration column (HiLoad 16/60 Superdex 200 prep grade, GE Healthcare). The protein sample was concentrated using Vivaspin 20 filter concentrators (VivaScience, USA), aliquoted into smaller fractions, flash frozen in liquid nitrogen and stored at −80°C.

### Pyrene-actin Polymerization Assays

Pyrene-actin polymerization assays were performed following the protocol described in previous studies (Miao et al., 2013; Sun et al., 2018). Rabbit skeletal muscle actin was purified from rabbit muscle acetone powder (Pel-Freez, USA). Pyrene-labeled actin was obtained from Cytoskeleton Inc. Monomeric actin at a concentration of 2 mM was mixed with 5% pyrene-labeled actin in the actin polymerization reactions. The fluorescence of pyrene was monitored using a fluorescence spectrophotometer (Cytation 5, BioTek, USA). The obtained data were analyzed and plotted using Origin software (Originlab Corporation, USA)

### RNA Sequencing (RNA-Seq) and analysis

Total RNA extraction was carried out using the GeneJET RNA Purification Kit (ThermoFisher Scientific, Waltham, MA) follwoing the manufacturers’ protocol. The purity of the extracted RNA was assessed using the Nanodrop 2000c Spectrophotometer (ThermoFisher Scientific, Waltham, MA). Subsequently, the RNA samples were sent to the RNA sequencing facility at Beijing Genomic Institute (Hong Kong) for cDNA library preparation and sequencing. For gene expression analysis, the numbers of reads uniquely mapped to the specific genes and the overall number of uniquely mapped reads in the sample were determined. The gene expression levels were calculated using the RPKM (Reads Per Kilobase per Million mapped reads) method.

### Image analysis

Post-acquisition image analyses were performed using either Fiji (Schindelin et al, 2012) or MATLAB (The MathWorks, Inc., Natick, MA). Kymographs were generated using the ‘Reslice’ tool. Depending on intensity of the fluorescent probes, an ‘average’ projection filters (average of 10 frames) and background subtractions may have been applied to enhance presentation. For consistency, the same processing was applied across all channels for kymographs generated from multi-color imaging. Additionally, the processing parameters were applied identically across different conditions that were directly compared to each other. Montages were generated using the ‘Make Montage’ tool. For intensity profiles, a region of interest with a size of 20 x 20 pixels was used for all movies. The intensity data points generated by Fiji were normalized and plotted using MATLAB. To visualize wave propagation, time projection images were created by merging three sequential frames at equal intervals and applying pseudocolors. To illustrate phase differences between proteins in wave propagation, multiple cycles of their intensity fluctuations were aligned. In the generated plots, solid lines represent the mean intensities, and shaded region represent the standard deviations of the intensities. To obtain the relative intensity, the maximum and minimum raw intensity values were determined and used to normalize the intensity value to a range of 0 −1. To assess the impact of FHDC1 knockdown on cell division, cells transfected with FHDC1 shRNA were imaged using DIC microscopy. The percentage of successful divisions was quantified by visually examining the images. Quantification of cell velocity was performed using the ‘Manual Tracking’ plugin in Fiji. The x and y coordinates of the center of the cell body was recorded and utilized to calculate the velocity of FHDC1 knockdown or control cells.

### Statistical analysis

All statistical analysis conducted using Prism 7 software (GraphPad Software, Inc, La Jolla, CA). To compare control and perturbed samples, unpaired two-tailed Student’s t-tests were performed. For multiple comparisons of frequencies in Fig 2D, a one-way ANOVA was performed. For multiple comparisons in Fig 4K, a one-way ANOVA followed by Sidak’s multiple comparison post hoc test was performed. Unless otherwise stated, error bars in all data shown represent mean ± S.E.M.

## Acknowledgements

We would like to express our gratitude to E. Feng and L. Cheung for their valuable technical assistance in this study. Additionally, we acknowledge the NTU Protein Production Platform (www.proteins.sg) for their support in the production of FHDC1 truncated variants. H.S was affiliated with the School of Biological Sciences, Nanyang Technological University, Singapore when the experiments were performed and is currently affiliated with the Department of Microbiology, Harvard Medical School, USA. X.C was affiliated with the Department of Cell Biology, Yale University School of Medicine, USA when the experiments were performed and is currently affiliated with the HiLIFE Institute of Biotechnology, University of Helsinki, Finland. R.R was affiliated with the Special Programme in Science at the National University of Singapore when the experiments were performed and is currently affiliated with the Department of Physics, Princeton University, USA. N.O was affiliated with the Special Programme in Science, National University of Singapore when the experiments were performed and is currently affiliated with the School of Life Sciences, University of Dundee, UK. This work is supported by the National Research Foundation (NRF) Singapore under its NRF Fellowship Program (M.W, NRF Award No. NRF-NRFF2011-09), Ministry of Education Academic Research Fund Tier 2 (M. W., 2015-T2-1-122) and Yale University startup grant (M. W). C.T is supported by a NUS Research Scholarship. M.S is supported by a MBI Scholarship. H.S is supported by MOE Tier 2 (MOE2016-T2-1-005S) to Y. M.

## Author contributions

C.T and M.S contributed equally to this paper. C.T, M.S, and M.W conceptualized and designed the experiments. C.T performed most of the experiments with contributions from S.M, X.C, R.R, N.O and A.L. C.T performed the data analysis. H.S. and Y.M contributed to the biochemical experiments. S.G contributed to the provisions of reagents. C.T, M.S and M.W interpreted results and contributed to the writing of the manuscript.

